# Acute pain drives different effects on local and global cortical excitability in motor and prefrontal areas: Insights into interregional and interpersonal differences in pain processing

**DOI:** 10.1101/2023.05.26.542414

**Authors:** Enrico De Martino, Adenauer Casali, Silvia Casarotto, Gabriel Hassan, Mario Rosanova, Thomas Graven-Nielsen, Daniel Ciampi de Andrade

**Affiliations:** Center for Neuroplasticity and Pain (CNAP), Department of Health Science and Technology, Faculty of Medicine, Aalborg University, Aalborg, Denmark; Institute of Science and Technology, Federal University of São Paulo, São Paulo, Brazil; Department of Biomedical and Clinical Sciences University of Milan, Milan, Italy; IRCCS Fondazione Don Carlo Gnocchi, Milan, Italy

**Keywords:** Transcranial magnetic stimulation, thermal pain, cortical activation, local and global cortical excitability, electroencephalogram, TMS-EEG

## Abstract

Pain-related depression of motor cortico-spinal excitability has been explored using transcranial magnetic stimulation (TMS)-based motor evoked potentials. Recently, TMS combined with concomitant high-density electroencephalography (TMS-EEG) enabled cortical excitability (CE) assessments in non-motor areas, offering novel insights into CE changes during pain states. Here, pain-related CE changes were explored in the primary motor cortex (M1) and dorsolateral prefrontal cortex (DLPFC). CE was recorded in 24 healthy participants before (Baseline), during painful heat (Acute Pain), and non-painful warm (Non-noxious warm) stimulation for eight minutes at the right forearm in a randomized sequence, followed by a pain-free stimulation measurement. Local CE was measured as peak-to-peak amplitude of the early latencies of the TMS-evoked potential (<120 ms) on each target. Furthermore, global-mean field power (GMFP) was used to measure global excitability. Relative to the Baseline, Acute Pain induced a decrease of −9.9±8.8% in the peak-to-peak amplitude in M1 and −10.2±7.4% in DFPFC, while no significant differences were found for Non-noxious warm (+0.6±8.0% in M1 and +3.4±7.2% in DLPFC; both P<0.05). A reduced GMFP of - 9.1±9.0% was only found in M1 during Acute Pain compared with Non-noxious warm (P=0.003). Participants with the largest reduction in local CE under Acute Pain showed a negative correlation between DLPFC and M1 local CE (r=-0.769; P=0.006). Acute experimental pain drove differential pain-related effects on local and global CE changes in motor and non-motor areas at a group level while also revealing different interindividual patterns of CE changes, which can be explored when designing personalized treatment plans.

**SUMMARY:** Cortical motor and prefrontal areas present reduced excitability during acute pain, but they occur in different patterns across individuals and present distinct impacts on global connectivity.

## INTRODUCTION

Acute experimental pain studies in healthy individuals have been widely used to investigate mechanisms underlying sensorimotor excitability, revealing an important effect of nociceptive system activation on sensorimotor plasticity [41]. By assessing somatosensory evoked potentials (SEPs) and motor-evoked potentials (MEPs), it has been demonstrated that acute pain inhibited sensorimotor excitability [8,47]. In contrast, intracortical inhibition was reported to be unchanged during acute pain [56] or decreased [23]. However, a major restriction of MEPs and SEPs is that they only allow the investigation of the local sensorimotor cortical excitability (CE) –added to the inherent contribution of spinal/peripheral excitability [52] - making it challenging to obtain a comprehensive picture of non-motor cortical areas responsible for the integration of affective, motivational, and evaluative aspects of pain. Additionally, it remains unknown to which extent CE changes in non-motor regions differ qualitatively from those in the motor cortex and to which degree these local CE changes engage global CE changes by alterations in functional connectivity related to acute pain. Understanding these mechanisms and relationships may shed light on the cortical dynamic changes related to acute pain and how they can be targeted more precisely by therapeutic interventions.

The development of TMS-compatible electroencephalography (TMS-EEG) technology has recently advanced the possibility of non-invasively probing local and global cortical excitability non-invasively in any cortical area targeted by transcranial magnetic stimulation (TMS) [27]. High-density EEG has been used to probe changes in cortical excitability following TMS of various brain targets [64] in physiological [42] and pathological [54] neurological conditions. These investigations rely on EEG recordings or local and global perturbation responses to single pulses of TMS to discrete cortical targets [49]. The TMS-evoked EEG potentials (TEPs) are reproducible waveforms with latencies before 300 ms generated by averaging EEG recording segments phase-locked to the TMS pulses [28] and provide a reliable measure of the local and global excitability of cortical circuits [4]. Therefore, TMS-EEG could be a valuable tool to study the motor and non-motor CE changes associated with pain, as well as the extent to which pain may modulate the engagement of global responses.

Understanding the differences in CE responses to pain is particularly relevant, considering these M1 and DLPFC targets are engaged in different steps in pain processing [25,45] and are also targets for therapeutic repetitive TMS [36]. Repetitive TMS to both M1 and DLPFC are known to be effective in some patients with acute and chronic pain but currently lack markers of therapeutic response [36], which could be revealed by exploring the connectivity pattern of these areas on an individual basis.

The current study compared for the first time the local CE responses in motor (M1) and non-motor (dorsolateral prefrontal cortex, DLPFC) areas induced by single-pulse TMS during experimentally induced acute heat pain and non-painful warm stimulation in healthy participants. Additionally, the global changes driven by local stimulation were assessed at both cortical targets as a measure of local-to-global connectivity.

## METHODS

### Participants

The current study was conducted at the Center for Neuroplasticity and Pain (CNAP), Aalborg University, Aalborg, Denmark, in October-December 2022. The study was performed according to the Helsinki Declaration, approved by the local ethics committee (Den Videnskabsetiske Komité for Region Nordjylland: N-20220018), and registered at ClinicalTrials.gov (NCT05566444) before the inclusion of participants. For the current repeated-measures analysis of variance (rmANOVA) design, a statistical power assessment using G*Power yielded a sample size estimate of 24 subjects based on the expected local and global CE changes (Power: 0.80, Alpha: 0.05, Effect size: 0.25, corresponding to η^2^_partial_ of 0.09 [14]). Twenty-four healthy volunteers were therefore recruited through online advertising, and written informed consent was signed before the commencement of the study. Participants were excluded if they suffered from acute and chronic pain, current and previous neurological, musculoskeletal, mental, and any other illnesses. Participants taking analgesic medication, or any other medication were also excluded. All participants were screened for contraindications to TMS [51], and completed the following questionnaires: Beck Depression Inventory [3], State-Trait Anxiety Inventory [61], Pain Catastrophizing Scale [62]; and Positive and Negative Affective Schedule [65] in order to assess normal mental health and the absence of catastrophic thinking related to pain.

### Experimental protocol

The experiment was conducted in a single experimental session, during which a total of eight TMS-EEG sessions were performed on two cortical areas: DLPFC and M1. Four different conditions were collected: Baseline, Acute pain, Non-painful warm, and Post (Figure 1B). To minimize potential bias, the order of cortical stimulation area (12 had M1 stimulation first) and Acute pain and Non-painful warm conditions (12 had Acute Pain first) were randomized across participants. The time interval between each condition was 5 minutes. A 30-minute interval was also included between M1 and DLPFC stimulation to allow time for finding the cortical spot, optimizing the EEG impendence, and ensuring that participants did not experience any discomfort or hear any clicking sounds during TMS stimulation.

**Figure 1:**
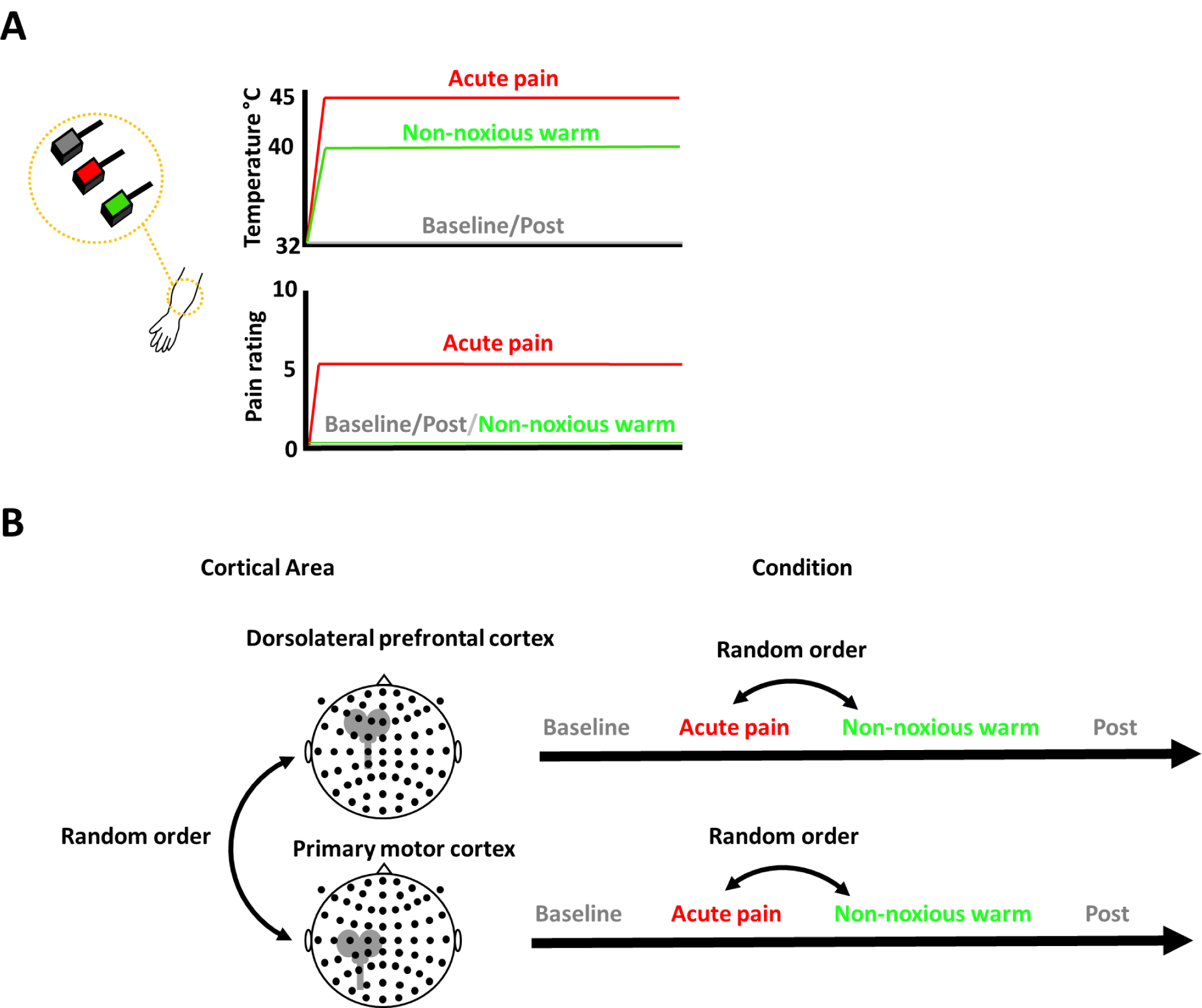
**A)** Baseline and Post (32°C), Non-painful warm (40.2±0.8°C) and Acute pain (45.2±0.8°C) temperatures were applied to the palmar region of the right forearm. Participants experienced pain intensity of around 5 using an 11-point scale during Acute Pain. **B)** Transcranial Magnetic Stimulation (TMS)-Electroencephalography was recorded in two Cortical Areas: The dorsolateral prefrontal cortex (DLPFC) and the primary motor cortex (M1). Four different conditions were recorded for each cortical area: Baseline, Acute pain, Non-painful warm and Post. The order of cortical stimulation area (DLPFC and M1) and Acute pain and Non-painful warm conditions were randomized.

### Thermal assessment and conditioned pain modulation

Individual pain thresholds were assessed using a thermal testing device (Medoc advanced medical systems, Haifa, Israel), with a thermode stimulator probe of 3 × 3 cm placed on the volar region of the right forearm (Figure 1A) and kept in place by medical tape and Velcro. Heat and cold pain thresholds were collected from a starting temperature of 32°C and increased (heat pain thresholds, HPT) or decreased (cold pain thresholds, CPT) by 1° C/s until the subject perceived a painful sensation occurred and pressed a stop button [48]. The interval between each measure was ∼30 seconds, and the mean of 3 successive measures was used.

Endogenous modulation of pain was assessed by conditioned pain modulation (CPM). According to CPM recommendations [66], participants immersed the left hand in a bucket of water and ice at ∼4° C for up to 60 seconds (cold pressor test) to feel moderately intense pain (∼7 on 10 on a 0-10 numerical rating scale, in which 0 indicates no pain and 10 indicates the most imaginable severe pain). Immediately after they withdrew their hand, HPT was reassessed. CPM was calculated as a difference from the HPT before conditioning (HPT during conditioning - HPT before conditioning).

### Acute pain and Non-painful warm stimulation

Participants were instructed to determine the intensity of the thermode stimulator probe required to cause moderately intense heat pain (Acute Pain) and an innocuous warm sensation (Non-painful Warm) during the TMS-EEG measurements (Figure 1A). Starting from the HPT, the temperature of the probe was increased in steps of 1 degree Celsius. Participants were asked to indicate when they perceived moderately intense heat pain, defined as a 5 out of 10 on a numerical rating scale, where 0 represents no pain and 10 represents the most severe pain imaginable. The value for Acute Pain conditions used during TMS-EEG data collection corresponded to temperatures between 44 and 46°C. A temperature between 39 and 41°C (below HPT) was used to induce the innocuous warm sensation. For Baseline and Post measurements, the thermode stimulator probe of 32°C (skin temperature) was used to avoid any thermal sensation (Figure 1A).

### TMS-EEG recording

Single-pulse TMS was delivered using a biphasic stimulator (Magstim Super Rapid^2^ Plus^1^, Magstim Co. Ltd, Dyfed, United Kingdom) and a figure-of-eight shaped coil (70 mm, Double Air Film Coil). To record TEPs, a TMS-compatible passive electrode cap with 63 electrodes (EASYCAP GmbH, Etterschlag, Germany) was placed according to the 10-5 system with the Cz electrode orientated to the vertex of the head. The ground electrode was placed halfway between the eyebrows. The electrode impedance was carefully monitored to ensure it remained below 5 kΩ. Raw recordings were sampled at a rate of 4800 Hz by a high-performance amplifier (g.HIamp EEG amplifier, g.tec-medical engineering GmbH, Schiedlberg, Austria) and online referenced to an additional forehead electrode (electrode 64). Two electrodes were also used to record the electrooculogram (EOG) laterally to the eyes (F9 and F10 electrodes). To mitigate the auditory responses to the click generated by the TMS coil from interfering with the TEPs, a TMS-click sound masking toolbox (TAAC, [53]) was utilized, and the participants wore noise-cancelling in-ear headphones (Shure SE215-CL-E Sound Isolating, Shure Incorporated, United States). To mitigate the somatosensory sensation produced by the TMS coil and EEG electrode movement artefacts, two net caps (GVB-geliMED GmbH, Ginsterweg Bad Segeberg, Germany) with a plastic stretch wrap handle film were applied to the EEG cap.

A navigated brain stimulation system (Brainsight TMS Neuronavigation, Rogue Research Inc., Montréal, Canada) was used to calibrate the participant’s head and TMS coil position using an optical-tracking system. To aid in localizing the M1 and DLPFC targets, a 3D reconstruction of the brain was generated by the navigated brain stimulation system using template MRI from Brainsight software (Rogue Research) scaled to the participant’s head. To identify the M1 target, the first dorsal interosseus (FDI) muscle over the left hemisphere was located in the proximity of the hand knob of the central sulcus [44], and the largest motor-evoked potential recorded by electromyographic electrodes was set as a “hot spot”. The rMT was established as the TMS intensity necessary to elicit MEPs greater than 50 μV in 5 of 10 trials delivering pulses at a minimum of 0.2 Hz, as measured from the FDI muscle electromyography [52]. The MEPs elicited in the FDI muscle were recorded with surface disposable silver/silver chloride adhesive electrodes (Ambu Neuroline 720, Ballerup, Denmark) parallel to the muscle fibers. A reference electrode was mounted on the ulnar styloid process. An intensity of 90% of the resting motor threshold (rMT) was applied for the M1 TEPs to avoid sensory-feedback contamination [21].

The DLPFC target was determined according to Mylius et al. 2013 in the middle frontal gyrus, and the stimulator intensity was set to 110% of the rMT of the FDI muscle [44]. To ensure the presence of a detectable TEP in both cortical targets, a real-time visualization tool (rt-TEP) was applied [11], which allowed slight adjustment of the TMS coil orientation across participants to minimize unwanted artefacts and ensured a minimum of 6 μV in the early peak-to-peak amplitude response in the average of 20 trials in the closest EEG electrode to DLPFC and M1 targets. The navigated brain stimulation system and rt-TEP were applied in real-time during the entire study to monitor the location of the TMS coil (within 3 mm of the cortical targets) and the highest signal-to-noise ratio in the EEG recordings. In each condition (Baseline, Acute Pain, Non-noxious Warm, and Post), about 160-180 pulses (∼8 min of TMS stimulation) were delivered with an interstimulus interval randomly jittered between 2600 and 3400 ms to avoid any significant reorganization/plasticity processes interfering with longitudinal TMS/EEG measurements [12]. During TMS stimulation, participants sat on an ergonomic chair with eyes open, looking at a fixation point on a wall. TEPs obtained from the stimulation of M1 and DLPFC waves were similar to those previously reported in clinical guidelines [64] (Figure 2).

**Figure 2:**
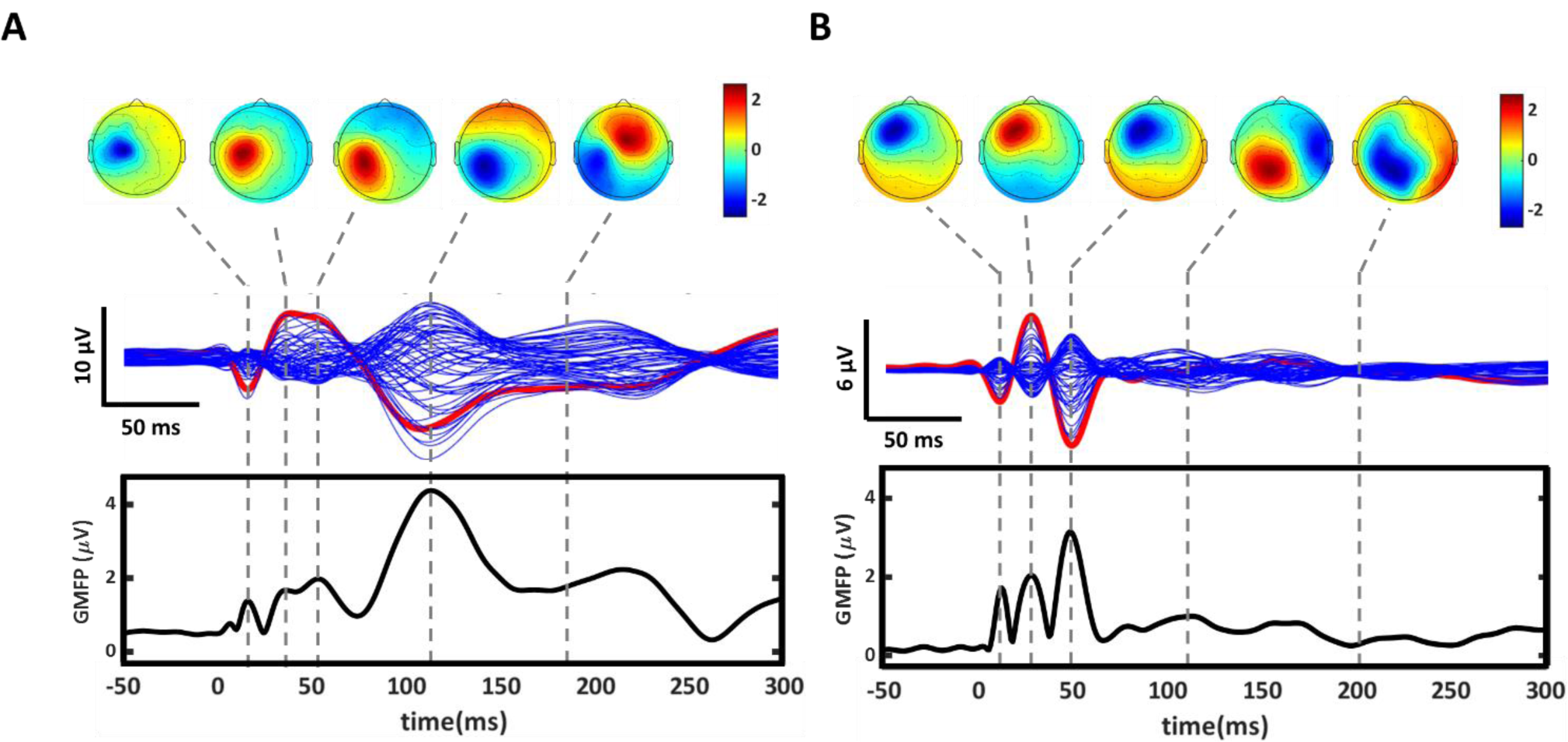
Sample data of TMS-evoked activity recorded with EEG following single pulse stimulation in a representative participant. **A)** The figure depicts the left primary motor cortex stimulation with the butterfly plot and topographical maps. The red line corresponds to the C3 electrode, and the blue lines correspond to the other 62 channels. The global mean field power (GMFP) of TMS-evoked potentials is shown below. **B)** The figure depicts the left dorsolateral prefrontal cortex with the butterfly plot and topographical maps. The red line corresponds to the F3 electrode, and the blue lines correspond to the other 62 channels. The global mean field power (GMFP) of TMS-evoked potentials is shown below.

### TMS-EEG data processing

Data were analyzed using Matlab R2019b (The MathWorks, Inc., Natick, MA, United States). All EEG recordings were split into epochs between ±800 ms around the TMS trigger. The TMS artefact was removed from all the EEG recordings replacing the recording between −2 and 6 ms from the TMS pulse with the time interval before the stimulation [15]. Bad epochs and channels containing noise, eye blinks, eye movements or muscle artefacts were visually inspected and rejected. The epochs were band-pass filtered (2±80 Hz, Butterworth, 3^rd^ order [21]) and down sampled to 1200 Hz. Channels were re-referenced to the average reference, and the four conditions (Baseline, Acute Pain, Non-painful warm and Post) were concatenated into a single trial. In this concatenated trial, independent component analysis (ICA, EEGLAB runica function [38]) was applied to remove any residual artefacts [50]. Then, epochs were segmented again in a time window of ±600 ms, and the concatenated trial was separated into the original four conditions (Baseline, Acute Pain, Non-painful warm and Post). Bad channels were interpolated using spherical splines [18].

Representing the local CE, the following parameters were extracted from the averaged artefact-free waveforms. The positive to negative peak-to-peak amplitude of the early component of TMS-evoked potentials (within 120 ms) were measured from the electrodes adjacent to the TMS stimulation site (C3, C1, Cp3, and CP1 electrodes were used for M1 cortical area and F3, F1, Fc3 and Fc1 electrodes were used for DLPFC cortical area) (Figure 3A and 4A). This early component is the highest reproducible across participants, and it is less affected by somatosensory or auditory responses [46]. This evoked component was comprised of a positive component between 25 and 50 ms (P30) followed by a negative deflection between 80 and 120 ms (N100) for M1 cortical area and a positive component between 15 and 30 ms (P25) followed by a negative deflection between 35 and 55 ms (N45) for DLPFC cortical area [64]. The positive and negative peaks and latency were extracted from each electrode and the measures from the four electrodes were averaged. Additionally, the peak-to-peak slope was calculated as the ratio between peak-to-peak amplitude and peak-to-peak interval (peak-to-peak amplitude/peak-to-peak interval). Based on animal studies, peak-to-peak amplitude and slope represent markers of synaptic strength [26]. Furthermore, the amplitudes and latencies of each peak-to-peak component were extracted to evaluate local cortical excitability according to clinical guidelines [64]. The percentage change from Baseline was calculated for the statistical analysis.

**Figure 3:**
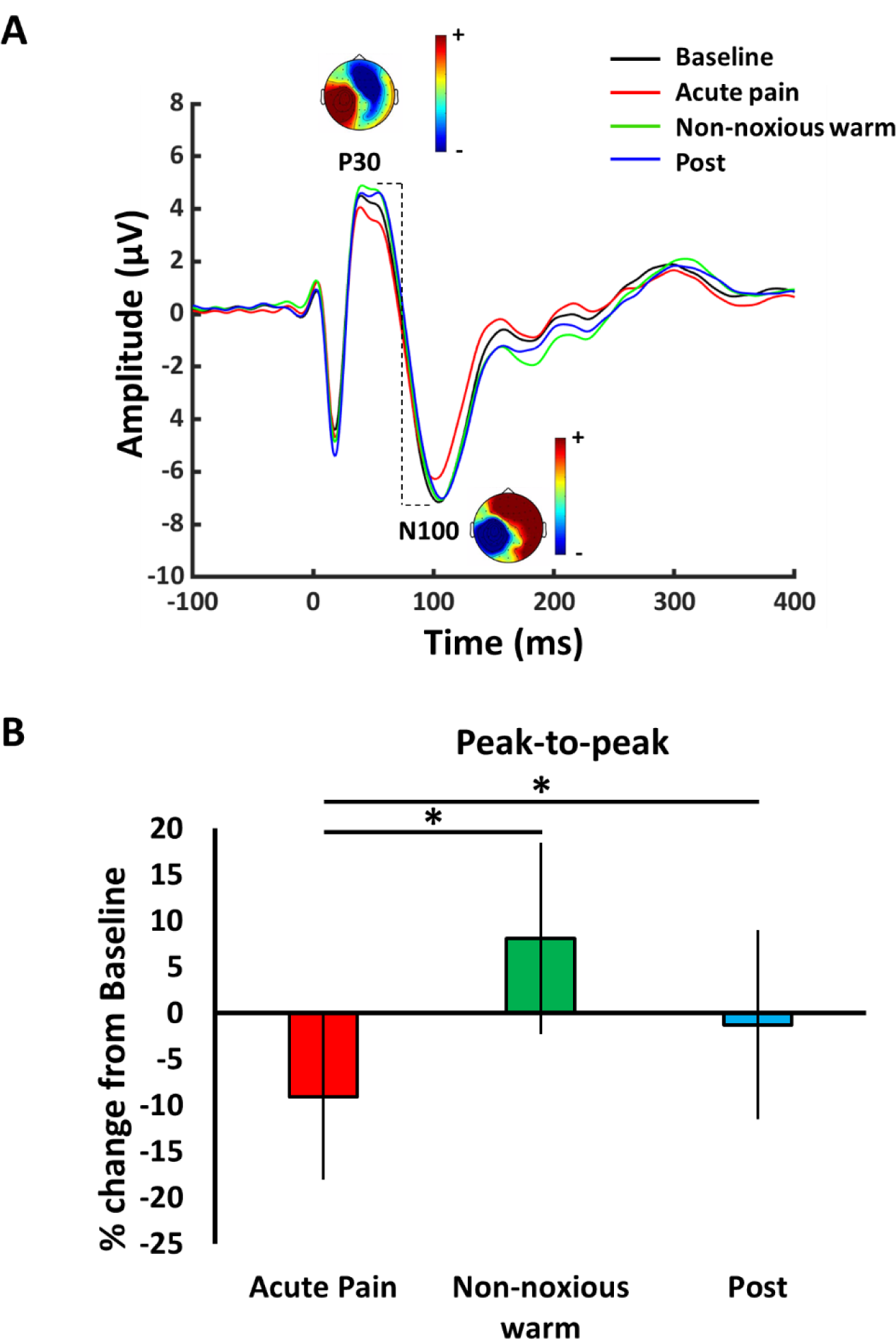
**A)** Grand average of TEPs (N = 24) during M1 stimulation at Baseline (black line), Acute pain (red line), Non-painful warm (green line) and Post (blue line). The largest peak-to-peak responses from the region close to the stimulation site were selected from each individual participant (selected channel for single participant: C3: n = 13; C1 n = 6; Cp3 n = 5). The topographical maps are shown for the 63 recorded channels next to the positive (P25) and negative (N45) peaks; **B)** Percentage changes from Baseline (mean and 95% confidence interval) are shown at Acute Pain, Non-painful warm and Post (Pairwise contrast * P <0.05).

**Figure 4:**
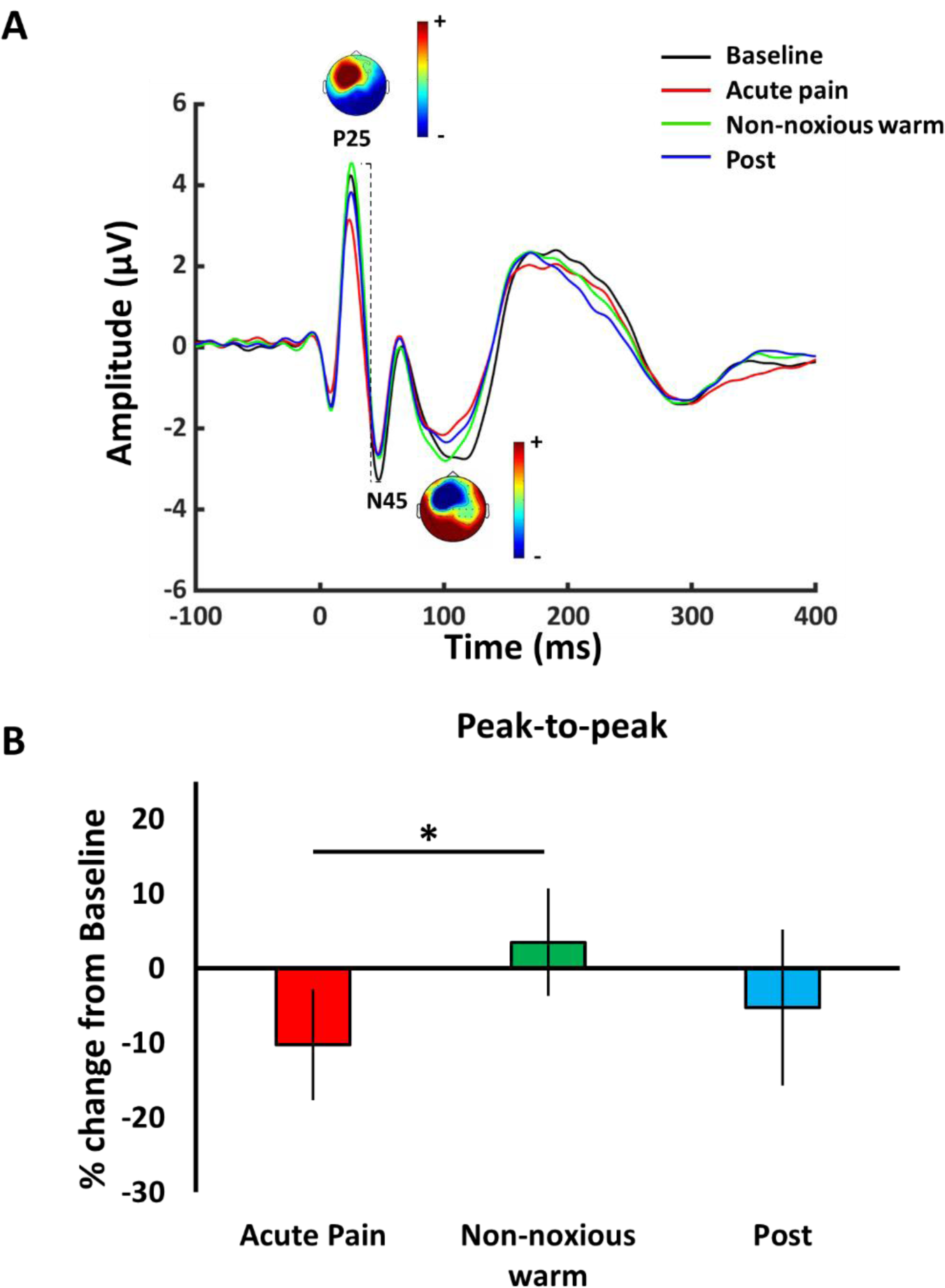
**A)** Grand average of TEPs (N = 24) during dorsolateral prefrontal cortex stimulation at Baseline (black line), Pain (red line), Warm (green line) and Post (blue line) conditions. The largest peak-to-peak responses from the region close to the stimulation site were selected from each individual participant (selected EEG channel for single participant: F1 n = 14; F3 n = 8; Fc1 n = 2). The topographical maps are shown for the 63 recorded channels next to the positive (P30) and negative (N100) peaks; **B)** Percentage changes from Baseline (mean and 95% confidence interval) are shown at Pain, Warm and Post conditions (*: significantly different from Pain, P <0.05).

For assessment of the global CE response, the global-mean field power (GMFP) was calculated as the root-mean-squared value of the signal across all electrodes in the 6-300 ms time interval after TMS stimulation (Figure 5). The GMFP estimates the amplitude dependence of the overall brain response [29] and allows the evaluation of the global cortical activity induced by the TMS stimulation as well as the indirect measurement of the connectivity degree between the stimulated target and distant brain areas [21]. In all cases, percentage change from baseline was used for comparisons between conditions.

**Figure 5:**
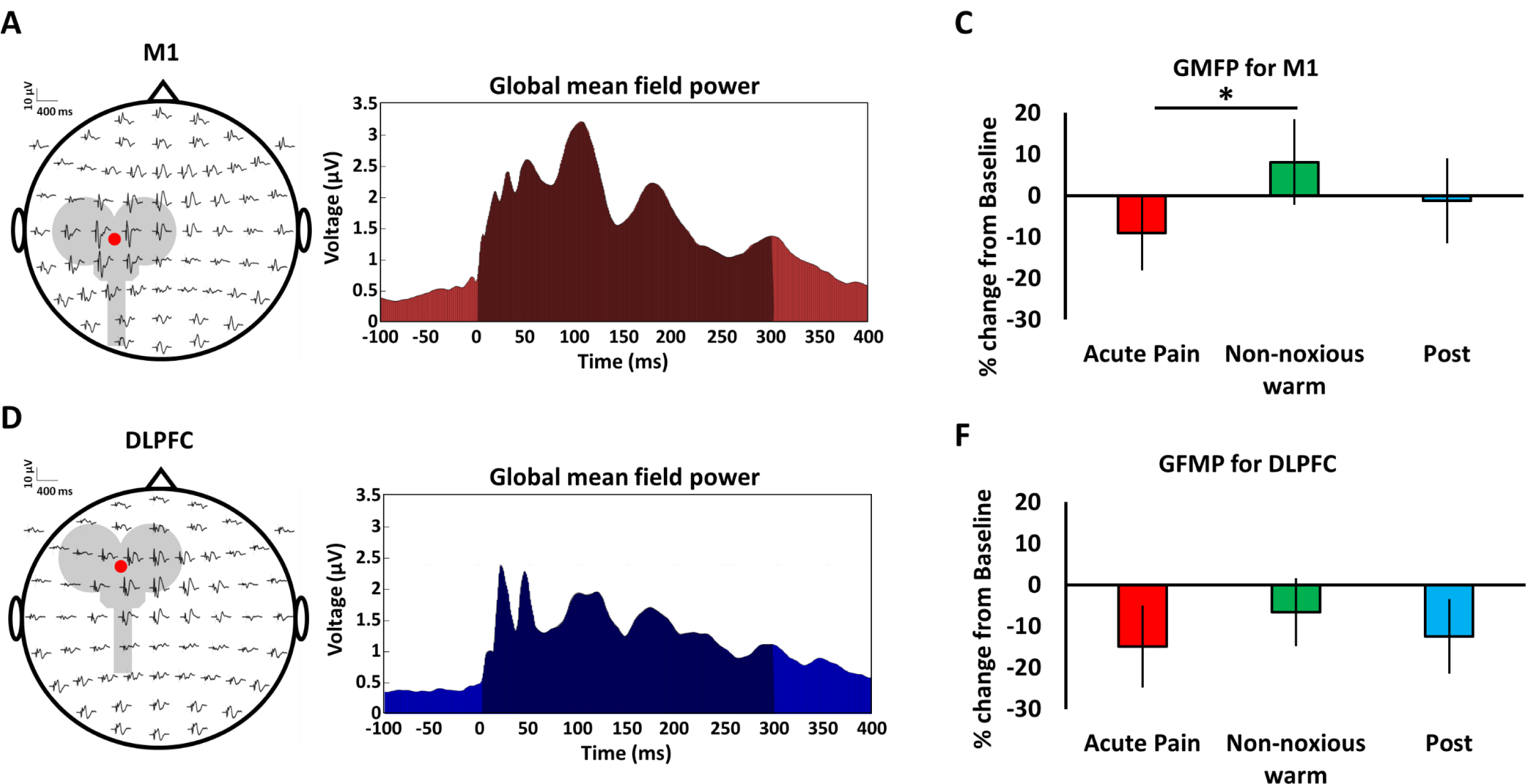
**A)** Grand average of the scalp maps (N = 24) presenting each individual electrode of TMS stimulation to M1. The red dot corresponds to the area of stimulation. **B)** Grand average of global-mean field power (Baseline; N = 24) from 63 EEG channels recorded in M1 at Baseline calculated as the root mean-squared value of the signal across all electrodes in the time interval of 6-300 ms after TMS stimulation (dark shaded area). **C:** Percentage changes from Baseline at Pain, Non-painful warm and Post (*: significantly different, P<0.05) for the M1 stimulation. **D)** Grand average of the scalp maps (N = 24) presenting each individual electrode of the TMS stimulation to DLPFC. The red dot corresponds to the area of stimulation. **E)** Grand average of global-mean field power (Baseline; N = 24) from 63 EEG channels recorded in DLPFC at Baseline. **F)** Percentage changes from Baseline at Acute Pain, Non-painful warm and Post conditions for the DLPFC stimulation.

### Statistical analysis

The Statistical Package for Social Sciences (SPSS, version 25; IBM, Chicago, United States) was used for statistical analysis. Results were presented as means and standard deviation, with a two-sided 5% significance level set for statistical significance. All parameters were assessed by visually examining histograms and Shapiro–Wilk tests. The Greenhouse−Geisser approach was used to correct violations of sphericity. Repeated measure ANOVAs with Condition (Acute Pain, Non-painful warm, and Post) as within-subject factors were used to analyze the percentage changes from the Baseline of peak-to-peak amplitude, slope, amplitudes, and latencies of the TEPs (for M1 P30 and N100 and for DLPFC P25 and N45) and GMFP in M1 and DLPFC. Effect sizes (partial eta squared: η^2^_partial_) were calculated in the statistical analysis [35]. Post hoc pairwise analyses were performed with Bonferroni-corrected multiple comparisons and corresponding 95% confidence intervals (95%CI) were generated when appropriate.

To determine whether different local cortical responses during Acute Pain differed among individuals, Pearson’s correlation analyses were conducted on the percentage changes from the Baseline of the peak-to-peak TEPs during Acute Pain in DLPFC and M1 cortical areas. The largest peak-to-peak TEP from the electrode closest to the stimulation site was selected from each individual participant (selected EEG channel for single participant for DLPFC: F1 n = 14; F3 n = 8; Fc1 n = 2; selected channel for single participant for M1: C3: n = 13; C1 n = 6; Cp3 n = 5). Furthermore, correlations were specifically performed on participants falling below the 1^st^ quartile in their peak-to-peak responses in DLPFC and M1 cortical areas. Pearson’s correlation analyses investigated the relationship between CPM, HTP, CPT and percentage changes of peak-to-peak TEP relative to Baseline. Finally, Pearson’s correlation analyses investigated the relationship between depression, anxiety, PANAS and pain catastrophizing and percentage changes of peak-to-peak TEP relative to Baseline.

## RESULTS

Data were successfully collected from 12 men and 12 women (Table 1). All psychological questionnaire-based parameters were within the normal ranges [3,61,62,65], as well as HPT, CPT, and CPM responses [17] (Table 1). During the TEPs recording, the participants rated the perceived pain as 5.1±1.3 during DLPFC stimulation and 5.0±1.2 during M1 stimulation on the NRS in the Acute Pain condition and zero in the other conditions.

**Table 1:** Mean (± standard deviation) of participant demographics, psychological questionnaires, and pain sensitivity.

| Variable |  |
| --- | --- |
| Age (years) | 27±5.5 |
| Height (cm) | 173±10.1 |
| Body mass (kg) | 70±14.2 |
| Depression (BDI-II) | 5.0±6.9 |
| State anxiety (STAI-S) | 42.6±6.1 |
| Trait anxiety (STAI-T) | 46.5±4.8 |
| Pain Catastrophizing scale | 8.7±7.9 |
| PANAS-negative | 13.3±3.0 |
| PANAS-positive | 25.0±5.4 |
| HPT ($^{\circ}$C) | 44.2±2.9 |
| CPT ($^{\circ}$C) | 13.1±9.0 |
| CPM effect ($^{\circ}$C) | 1.4±1.2 |

BDI-II, beck depression inventory; STAI, state-trait anxiety inventory; PANAS, positive and negative affective schedule; HPT, heat pain thresholds; CPT, cold pain thresholds; CPM, conditioned pain modulation.

### M1 local excitability and pain

The grand averages of TEPs obtained from the TMS stimulation of the left M1 cortical area during the four conditions are shown in Figure 3A. The intensity used on M1 during the EEG recording was 58.0±9.5% (rMT of the FDI muscle was 64.3±9.5%). The average number of artefact-free epochs used to calculate the TEPs was 150±11 at Baseline, 154±13 at Acute Pain, 150±11 at Non-noxious warm, and 149±12 at Post condition.

The rmANOVA showed a significant effect of Condition for the peak-to-peak amplitude (Figure 3B; F_(2,46)_ = 5.01; P = 0.010; η^2^_partial_ = 0.19). Post-hoc tests revealed a difference between Acute Pain (−9.9±8.3%) and Non-noxious warm (0.6±8.0%, P = 0.037) and between Acute Pain and Post condition (0.6±5.3%, P = 0.036). Similarly, the rmANOVA showed a significant effect of Condition for the peak-to-peak slope (F_(2,46)_ = 5.42; P = 0.008; η^2^_partial_ = 0.19). Post-hoc testing revealed a difference between Acute Pain (−10.3±8.4%) and Non-noxious warm (0.6±5.8%, P = 0.034) and between Acute Pain and Post condition (0.7±4.3%, P = 0.049).

Analyzing the P30 and N100 separately, the rmANOVA only revealed a significant effect of Condition for the P30 amplitude (F_(2,46)_ = 7.01; P = 0.002; η^2^_partial_ = 0.23), but no effect for the N100 (F_(2,46)_ = 2.45; P = 0.098; η^2^_partial_ = 0.10). Post-hoc tests revealed a difference in P30 amplitude between Acute Pain (−11.9±10.4%) and Non-noxious warm (6.3±11.0%, P = 0.003) and between Acute Pain and Post condition (6.4±11.0%, P = 0.006). No significant difference was found for P30 and N100 latencies (both F_(2,46)_ < 3; P > 0.05; η^2^_partial_ ≤ 0.10). The individual data for M1 stimulation and non-normalized parameters are reported in the supplementary material (Supplementary Figures 1 and 2, Tables 1 and 2).

### DLPFC local excitability and pain

The grand averages of TEPs obtained from the TMS stimulation of the left DLPFC cortical area are shown in Figure 4A. The intensity used on DLPFC during the EEG recording was 70.7 ± 10.5% of TMS stimulator output. The average number of artefact-free epochs used to calculate the TEPs for DLPFC was 145 ± 18 at Baseline, 151 ± 18 at Acute Pain, 150 ± 18 at non-noxious warm, and 149 ± 23 at Post condition. The rmANOVA showed a significant effect of Condition for the peak-to-peak amplitude (F_(2,46)_ = 4.82; P = 0.013; η^2^_partial_ = 0.17). Post-hoc test revealed a difference between Acute Pain (−10.2±7.4%) and Non-noxious warm (3.5±7.2%, P = 0.030), but no significant difference was found between Acute Pain and Post condition (−5.2±10.4%, P = 0.540, Figure 4B). The rmANOVA also showed a significant effect of Condition for the peak-to-peak slope (F_(2,46)_ = 5.94; P = 0.005; η^2^_partial_ = 0.21). Post-hoc tests revealed a difference between Acute Pain (−8.9±8.7%) and Non-noxious warm (6.9±12.8%, P = 0.026), and between Acute Pain and Post condition (8.7±13.1%, P = 0.018).

When the amplitude of P25 and N45 was separately analyzed, the rmANOVA revealed a significant effect of Condition for P25 (F_(2,46)_ = 4.48; P = 0.017; η^2^_partial_ = 0.16), but no effect for N45 (F_(2,46)_ = 0.14; P = 0.871; η^2^_partial_ = 0.01). Post-hoc tests for the P25 showed a difference between Acute Pain (−16.7±8.9%) and Non-noxious warm (0.3±8.7%) (P = 0.001), but no difference was found between Acute Pain and Post (−13.8±15.2%, P = 0.969). No significant difference was found for P25 and N45 latencies (both F_(2,46)_ < 3; P > 0.05; η^2^_partial_ ≤ 0.12). The individual data and non-normalized parameters are reported in the supplementary material (Supplementary Figures 1 and 2, Tables 1 and 3).

### Correlations of local excitability changes during pain

No significant correlation was found between changes in peak-to-peak TEP amplitude under Acute Pain between M1 and the DLPFC (r = −0.131; P = 0.543, Figure 6A). The visual inspection of individual patient data showed that most healthy participants had reduced peak-to-peak amplitudes in either M1 or the DLPFC. A subsequent correlation analysis of the participants with intense peak-to-peak reduction (below the 1^st^ quartile (N=11)) revealed a negative correlation between DLPFC and M1 change in amplitude during acute pain (r = −0.769; P = 0.006) (Figure 6B), indicating that in almost 50% of healthy individuals, Acute Pain induced marked changes in local CE in one of the two targets, while only one participant had major amplitude decreases concomitantly in both targets (Figure 6C).

**Figure 6:**
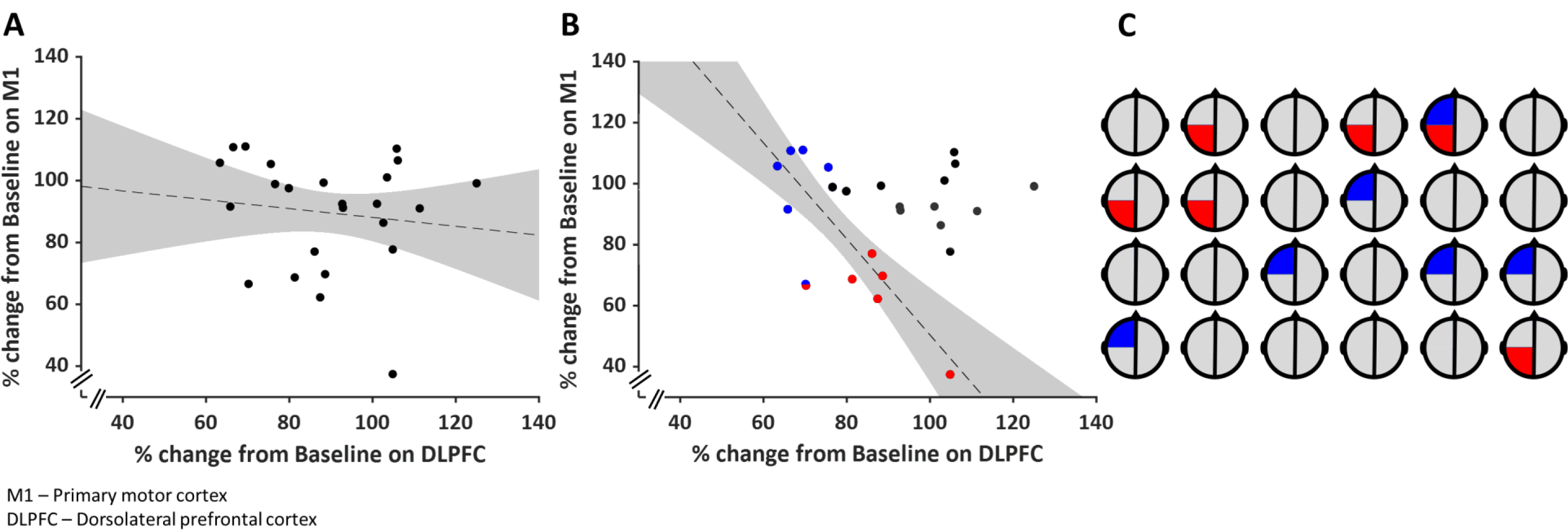
Correlation between the peak-to-peak responses from M1 and DLPFC stimulation expressed as a percentage of the Baseline from Acute pain. **A)** Correlation between M1 and DLPFC stimulation in 24 participants (grey shaded area indicates 95% confidence intervals). **B)** Correlation between the peak-to-peak responses from M1 and DLPFC stimulation in the participants who exhibited a falling below the first quartile in their responses. The blue dots represent participants showing a falling below the first quartile in DLPFC stimulation, while the red dots represent participants showing a falling below the first quartile in M1 stimulation. **C)** The figure presents 24 stylized heads, each representing an individual participant (heads marked with a red quarter denote participants who exhibit reductions below the 1^st^ percentile in peak-to-peak amplitude in M1, and a blue indicates participants with reductions below the 1^st^ percentile in peak-to-peak amplitude in DLPFC.

No correlations were found between CPM, HTP, CPT and the percentage change during Acute Pain relative to the Baseline in peak-to-peak amplitude in DLPCF (all P > 0.05) and M1 (all P > 0.05) cortical area. No correlations were found between questionnaires and the percentage change during Acute Pain relative to the Baseline in peak-to-peak amplitude in DLPCF and M1 (all P > 0.05).

### Global responses during pain

For the M1, the rmANOVA of the global mean field power showed a significant Condition effect (F_(2,46)_ = 5.48; P = 0.007; η^2^_partial_ = 0.19). Post-hoc tests revealed a difference between Acute Pain (−9.1±8.9%) and Non-noxious warm (8.1±10.4%, P = 0.003), but no significant difference was found between Acute Pain and Post condition (−1.3±10.3%, P = 0.301, Figure 5C). No significant effect of Condition was found (F_(2,46)_ = 1.46; P = 0.242; η^2^_partial_ = 0.06) for DLPFC (Figure 5F). The individual data and non-normalized GMFP for DLPFC and M1 are reported in the supplementary material (Supplementary Figure 2 and Table 1).

## DISCUSSION

The present results provided novel insight into local and global CE changes in motor and non-motor areas in acute experimental pain. The findings revealed that the decrease in CE during acute pain in M1 and the DLPFC was primarily influenced by the first positive component of the TMS-evoked EEG response and that participants with the greatest reduction in peak-to-peak percentage change in the DLPFC were not the same presenting the largest decreases in the sensorimotor cortex. Only M1 appeared to be engaged in a significant global reduction in cortical excitability during acute pain, suggesting differential local-to-global dynamics between these two areas during acute experimental pain.

### Decrease in local cortical excitability

Previous attempts to measure CE during acute pain were performed by assessing MEP recorded from the upper limb and face muscles [8,47]. Since CE obtained by MEPs represents a cumulative measure of excitability across cortico-cortical, cortico-motoneuronal, and spinal motoneuron synapses, alterations at spinal and peripheral levels could potentially contribute to the MEPs reduction in previous studies [8,47]. In the current study, subthreshold intensities were used to stimulate M1 during acute experimental pain. The reduction of the local CE responses indicates a localized reduction in the cortico-cortical excitability at the motor cortex level or in its local surrounding cortical-subcortical networks [5]. Notably, the current study showed a specific reduction of the P30 amplitude (the first positive component of the M1 response to TMS), which was previously correlated to MEPs response, at least in threshold intensities [39]. The origin of P30 is only partially known, and it has been suggested to reflect the activity of the ipsilateral premotor [20] and supplementary motor areas [37] since 5Hz rTMS over the hand area of M1 increases this peak [20] and has a metabolic effect on premotor and supplementary motor areas [60]. Furthermore, an increased P30 amplitude has been described in patients with progressive myoclonus epilepsy [29], which supports the possibility that the P30 has a motor/premotor origin.

Finally, the administration of the voltage-gated sodium channel blocker carbamazepine can suppress P30 in healthy participants, indicating a reduction of motor CE [16]. Together with previous studies, the current results suggest that nociceptive inputs, but not warm, reduce motor excitability, as well as the surrounding motor areas, as part of the central plastic changes related to pain.

Inhibitory effects were also found when left DLPFC was probed during acute pain (reduction in the peak-to-peak amplitude and P25 amplitude), pointing towards an analogous local reduction of CE in the prefrontal cortex. In previous studies, peak-to-peak amplitude and slope in the local electrical response of neurons have been linked to a buildup of CE and synaptic strength in the left frontal cortex [10,26]. This association was clear when, after a night of sleep deprivation, peak-to-peak amplitude and slope increased significantly in healthy participants, being subsequently rebalanced after a night of undisturbed sleep recovery [26]. The increased CE was confirmed in vitro and in vivo animal experiments during wakefulness [9]. Additionally, electroconvulsive therapy increased the peak-to-peak amplitude and slope of the electrical response of neurons in the left frontal cortex excitability in patients affected by severe major depression [10], suggesting normalized frontal CE in patients with dysfunctional limbic-cortical pathways [43]. Notably, in the current study, P25 (the first positive component of the TEP after probing the DLPFC) was affected by acute pain and not by warm. This positive component from DLPFC stimulation has been related to frontal CE since intermittent theta-burst stimulation to DLPFC decreased the amplitude of this TEP component in healthy individuals [19]. In addition to mood disorders, the left DLPFC is also known to play a critical role in pain appraisal and detection [57]. According to neuroimaging studies, acute and chronic pain conditions are commonly associated with reduced left DLPFC structure and function [2], reflecting probably a hypo-metabolic state [58]. By contrast, pain-relief interventions, such as therapeutic high-frequency left DLPFC repetitive TMS, can reverse this reduced function, and, have been applied as a modulatory strategy to experimentally induced pain [40,63], post-surgical pain [6,7] and chronic pain [59]. Together with previous studies, the current findings indicate that nociceptive synaptic transmission modulates the left prefrontal activity towards reduced CE compared to the warm control condition.

A key finding of the current study was that decreases in M1 and DLPFC local CE were significant and present at similar magnitudes but did not correlate. After individual assessment of each participant’s response profile, it was clear that those exhibiting the most significant decrease in peak-to-peak decrease at DLPFC were distinct from those with the largest reduction at M1 stimulation. In fact, about 50% of participants were either polarized to M1 or to DLPFC as the main reduction in excitability site and in this subgroup comprising more than half of the participants, a high negative correlation was found between M1 and DLPFC changes during acute pain. This finding suggests a non-uniform interindividual cortical response to acute pain in healthy people, with some individuals presenting changes in a cognitive-evaluative network hub (DLPFC), and others in a sensorimotor one (M1). This is a novel finding that may help reveal the grounds for the interindividual differences in pain perception and treatment response also seen in patients [24]. Current pain treatment strategies primarily follow a “one-size-fits-all” approach, wherein patients receive the same treatment regardless of their individual characteristics. As a result, a significant number of patients remain resistant to treatment, with up to 50% of chronic pain sufferers experiencing symptoms despite optimal medical care [1]. TMS-EEG may present a promising non-invasive electrophysiological method to investigate cortical function during pain [22].

### Globally, not all targets are equal

The current study found a decrease in global CE during acute pain only when probing M1, but this was not found significant for the DLPFC. M1 and the DLPFC have different structural and functional connectivity profiles and are hubs in different brain networks. These differences can be appraised in the dynamics of M1 activity propagations after a probing pulse to M1: after the engagement of the stimulation target, activity spreads to more postero-lateral locations, via corticocortical volleys from M1 to the somatosensory areas, and to the opposite hemisphere, via the corpus callosum or subcortical pathways [33]. Differently, the left DLPFC TMS activates the site of stimulation as well as the opposite prefrontal cortex [30,34]. Previous research has also shown that TMS delivered over M1 at rest results in a larger global response compared to DLPFC when using the same TMS intensity [31,34]. This difference is even more pronounced when TMS to M1 is delivered above the rMT [31] due to peripheral nerve stimulation at suprathreshold intensities [32]. In the current study, higher TMS intensity was used to stimulate the DLPFC (110% of the FDI rMT) compared to M1 (90% of the FDI rMT) to produce a clear and reproducible cortical response for DLPFC stimulation. Additionally, baseline normalizations were used so that the intrinsic baseline activation threshold between targets would be compensated for. And still, only M1 showed significant global CE reduction during acute pain compared to non-noxious stimulation. This is an original finding that highlights qualitative changes in the extent to which M1 connectivity is affected by acute pain.

### Limitations

The results of the current study should be interpreted in consideration of its limitations. First, the CE responses induced by TMS can be contaminated by auditory and somatosensory responses [4,13]. To minimize this, we included control conditions with warm or no stimulation and compared differences based on changes from baseline values. More importantly, only earlier and local peaks (below 120 ms) were assessed, as auditory and somatosensory responses mainly impact the 100-200 ms range despite optimal measures to minimize them [46]. For the GMPF, the time interval analyzed was from 6 to 300 ms after the TMS stimulation to assess the entire CE and the excitability of remote cortical regions. Since only GMFP from M1 was reduced during acute pain, it is possible to exclude the contribution of potential non-specific effects, such as decreases in arousal, attention, and reduced auditory or somatosensory responses. Secondly, this study did not assess the saliency of the non-noxious warm stimulus, which might affect the results. The warm condition was applied as a control condition to provide similar sensory inputs to the forearm, which could interfere with the cortical response and were not painful. Despite this measure, acute pain and non-noxious stimuli engage saliency systems differently. While it may be argued that activation of the saliency system is an intrinsic component of the pain experience, it can be controlled by matching saliency from non-painful stimuli, which was not done here. Finally, early peaks in the remote regions of the cortex (i.e., right M1 or the right DLPFC [64]) were not analyzed since the main aim of this study was to assess local CE responses during acute pain. Nevertheless, non-motor TEP responses during acute pain could be relevant as contralateral motor plastic changes to a painful stimulus were described using traditional MEP measurements [55].

