## Supplementary tables and figures for "Acute pain drives different effects on local and global cortical excitability in motor and prefrontal areas: Insights into interregional and interpersonal differences in pain processing"

### SUPPLEMENTARY MATERIAL

#### *Supplementary figure*

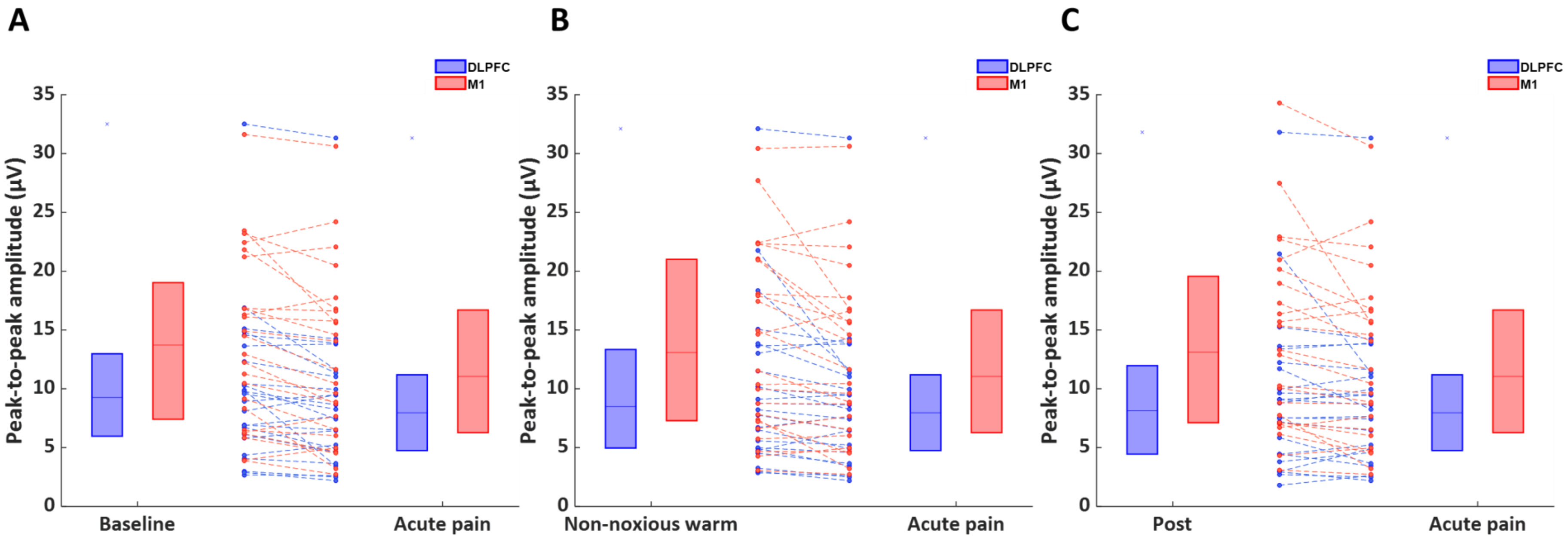


**Supplementary Figure 1**: Changes in peak-to-peak amplitude resulting from stimulation of the DLPFC and M1. The blue box plot (median and first and third quartiles) and blue dots (data from individual participants) represent DLPFC stimulation, whereas the red box plot (median and first and third quartiles) and red dots (data from individual participants) represent primary motor cortex (M1) stimulation. **A)** Comparison between Baseline and Acute Pain; **B)** Comparison between Non-noxious warm and Acute Pain; **C)** Comparison between Post and Acute Pain.

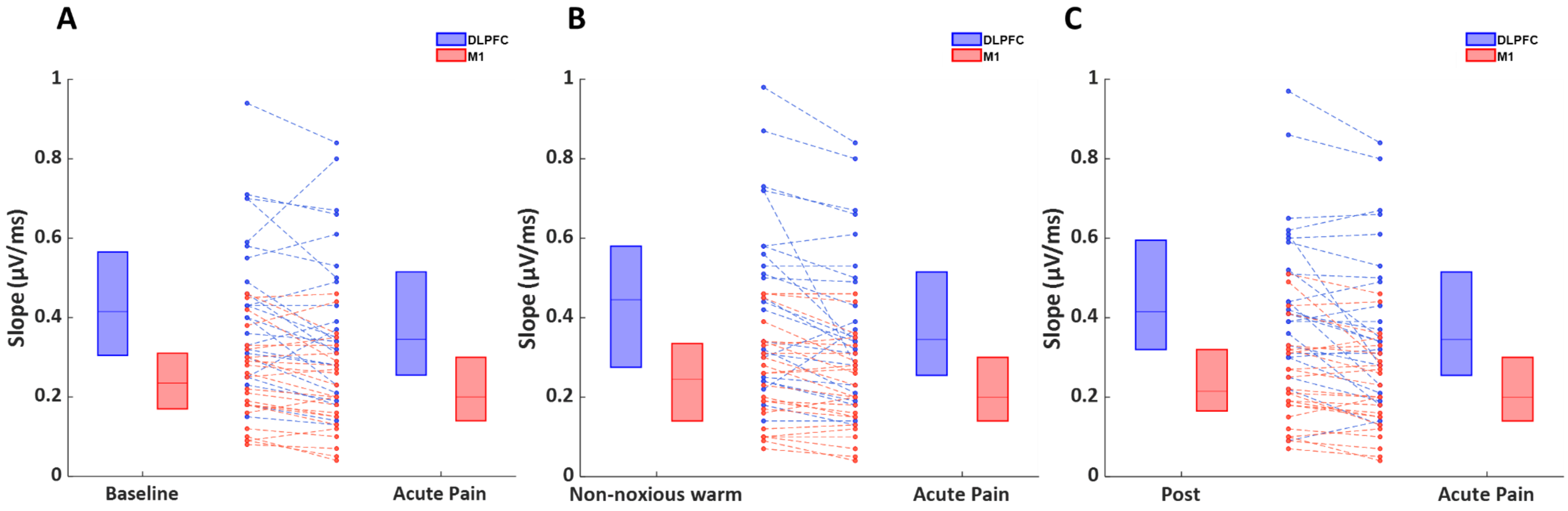


**Supplementary Figure 2:** Changes in peak-to-peak slope resulting from stimulation of the DLPFC and M1. The blue box plot and blue dots (data from individual participants) represent DLPFC stimulation, whereas the red box plot and red dots (data from individual participants) represent M1 stimulation. **A)** Comparison between Baseline and Acute Pain; **B)** Comparison between Non-noxious warm and Acute Pain; **C)** Comparison between Post and Acute Pain.

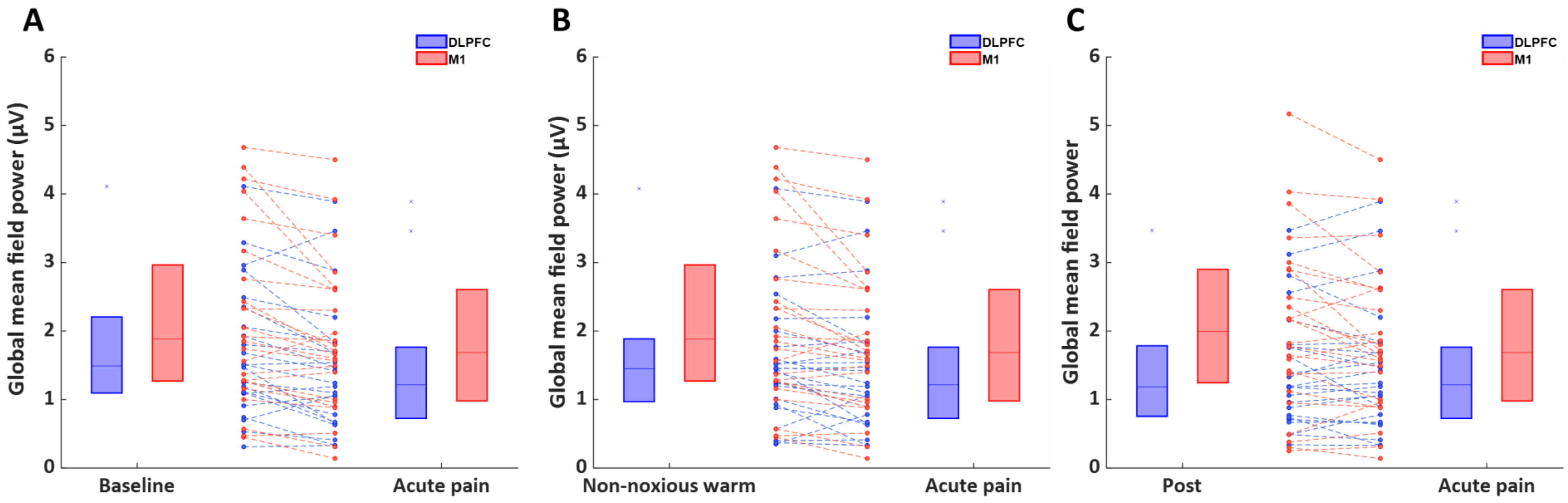


**Supplementary Figure 3:** Changes in global mean field power (GMFP) resulting from stimulation of the DLPFC and M1. The blue box plot (median and first and third quartiles) and blue dots (data from individual participants) represent DLPFC stimulation, whereas the red box plot (median and first and third quartiles) and red dots (data from individual participants) represent M1 stimulation. **A)** Comparison between Baseline and Acute Pain; **B)** Comparison between Non-noxious warm and Acute Pain; **C)** Comparison between Post and Acute Pain.

### *Supplementary tables*

**Supplementary table 1.** Mean ± standard deviation of peak-to-peak amplitude, peak-to-peak slope, and global mean field power in DLPFC and M1 during Baseline, Acute pain, Non-noxious warm and Post conditions.

| Variable | Cortical Area | Condition | | | |
|---|---|---|---|---|---|
| | | Baseline | Acute pain | Non-noxious warm | Post |
| Peak-to-peak amplitude (µV) | DLPFC | 9.84±6.36 | 8.76±6.06 | 10.02±6.83 | 9.35±6.68 |
| | M1 | 13.76±7.02 | 12.67±7.58 | 14.06±6.82 | 14.03±7.94 |
| Peak-to- peak Slope (µV/ms) | DLPFC | 0.44±0.19 | 0.40±0.20 | 0.46±0.22 | 0.46±0.20 |
| | M1 | 0.24±0.11 | 0.22±0.11 | 0.25±0.12 | 0.25±0.12 |
| Global mean field power (µV) | DLPFC | 1.68±0.94 | 1.43±0.93 | 1.54±0.90 | 1.44±0.86 |
| | M1 | 2.01±1.06 | 1.86±1.10 | 2.16±1.27 | 2.07±1.26 |

DLPFC, dorsolateral prefrontal cortex; M1, primary motor cortex.

**Supplementary table 2.** Mean ± standard deviation of the amplitude of the TEPs in M1 stimulation during Baseline, Acute pain, Non-noxious warm, and Post conditions.

| Variable | Condition | | | |
|---|---|---|---|---|
| | Baseline | Acute pain | Non-noxious warm | Post |
| P30 amplitude (µV) | 6.34±3.90 | 5.75±3.99 | 6.78±4.61 | 6.79±4.63 |
| N100 amplitude (µV) | -7.41±3.85 | -6.92±4.08 | -7.29±3.89 | -7.25±4.13 |
| P30 latency (ms) | 42.2±11.8 | 40.4±12.4 | 42.+±10.6 | 42.0±12.1 |
| N100 latency (ms) | 101.1±23.3 | 99.5±23.4 | 100.7±22.4 | 101.6±23.1 |

**Supplementary table 3.** Mean ± standard deviation of the amplitude of the TEPs in DLPFC stimulation during Baseline, Acute pain, Non-noxious warm, and Post conditions.

| Variable | Condition | | | |
|---|---|---|---|---|
| | Baseline | Acute pain | Non-noxious warm | Post |
| P25 amplitude (µV) | 5.34±3.58 | 4.34±2.90 | 5.56±4.30 | 4.78±4.16 |
| N45 amplitude (µV) | -4.50±3.76 | -4.42±3.86 | -4.60±3.63 | -4.57±2.98 |
| P25 latency (ms) | 22.19±2.90 | 21.11±2.51 | 22.67±3.63 | 21.88±3.89 |
| N45 latency (ms) | 43.92±6.28 | 42.22±7.58 | 44.34±7.28 | 41.58±8.40 |
